# Efficient Generation of Human Pluripotent Stem Cells from Frozen Cord Tissue via Chemical Reprogramming

**DOI:** 10.1101/2024.04.10.588154

**Authors:** Coneria Nansubuga, Donna K. Mahnke, Siqi Li, Abigail Multerer, Jake Minx, Bradley Miller, Mengcheng Shen, Andreas Beyer, Lu Han, Joy Lincoln, Chun Liu

## Abstract

Chemical reprogramming presents an innovative approach for generating induced pluripotent stem cells (iPSCs), bypassing the genetic instability and safe concern associated with viral vector approach. We describe a novel, efficient chemical method for reprogramming human umbilical cord tissue-derived mesenchymal stem cells (MSCs) into induced pluripotent stem cells (iPSCs). Compared to previous sources like adipose tissue and skin, frozen umbilical cord tissue offers an abundant, non-invasive, long-term storable, and ethically sound cell source. Our findings not only showcase the feasibility and safety of utilizing chemical reprogramming on cells from frozen umbilical cords but also underscore its potential in regenerative medicine, especially for developing safer and more effective therapies for cardiovascular diseases.

## INTRODUCTION

Chemical reprogramming offers a compelling alternative to viral methods for generating induced pluripotent stem cells (iPSCs), a cornerstone in regenerative medicine and biomedical research^1^. Unlike viral reprogramming, which introduces genetic materials through vectors with potential for insertional mutagenesis, chemical reprogramming employs small molecules to overpass the genetic modifications, thus minimizing risks of genetic instability and enhancing safety profiles ^1,2^. This approach not only mitigates concerns related to genomic alterations but also offers a more controllable and reversible mechanism, critical for clinical applications. Furthermore, the scalability and cost-effectiveness of chemical methods surpass those of viral techniques, presenting a more feasible pathway for widespread therapeutic use and personalized medicine^3,4^. By addressing the limitations associated with viral vectors, chemical reprogramming emerges as a promising avenue, potentially revolutionizing the generation of iPSCs for the study and treatment of cardiovascular diseases and beyond.

Current human chemical reprogramming method was primarily conducted on human mesenchymal stem cells (MSCs) from adipose or dermis tissues^1,3^. However, umbilical cord tissue emerges as a superior alternative for several reasons: it is an abundant source of MSCs ^5^; its frozen form ensures both sustainability and easy accessibility of cells without necessitating invasive procurement methods like blood PBMCs^6^; MSCs from umbilical cord tissue exhibit a higher proliferation rate, potentially increasing the efficiency of reprogramming processes ^7^. Furthermore, given that iPSC reprogramming efficiency is much higher when using young cells, umbilical cords represent a superior cell source compared to blood cells from older donors ^8^. Here, for the first time, we successfully employ chemical reprogramming on MSCs isolated from frozen umbilical cord tissues. It not only enriches the toolkit of regenerative medicine but also paves the way for innovative therapeutic strategies that are safer, more efficient, and ethically sound.

## Main Text

### Isolate MSCs from frozen cord tissues

Umbilical cord tissues were initially processed and cryogenically stored at the Cord Blood Registry in Tucson, AZ. The process began with cutting the tissue into 1 cm segments within a clean petri dish. These segments were further cut into halves and placed into a 50 mL tube filled with PBS for rinsing. Subsequently, the halves were transferred to another sterile dish, and a 4-mm biopsy punch was used to obtain tissue pieces with a uniform size. These samples were then allocated into 5 mL cryotubes, each filled with 2 mL cryopreservation solution containing approximately 2-2.5 g of tissue. Initially, the samples were frozen at -80°C, and after 18-24 hours, they were transferred to liquid nitrogen (LN2) tank for long-term preservation. To isolate MSCs for iPSC reprogramming, the frozen tissues were quickly thawed in a 37°C water bath on site (Figure 1A). Tissue segments were then placed on a 5 x 5 grid on a 10 cm culture dish coated with a MSC attachment-promoting substrate. To ensure firm attachment, tissues were left to dry for 10 mins before gradually adding 15 mL of complete MSC media. After 7-8 days of incubation, fibroblast-like cells migrating out from the periphery of umbilical cord tissues were observed. Then original tissue segments were carefully removed and cells were expanded in fresh MSC media. MSCs became confluent between day 14 and day 16, at which point cells were further expanded for iPSC reprogramming or cryopreservation. The MSCs from frozen umbilical cord tissues are referred to ucMSCs.

**Figure 1.**
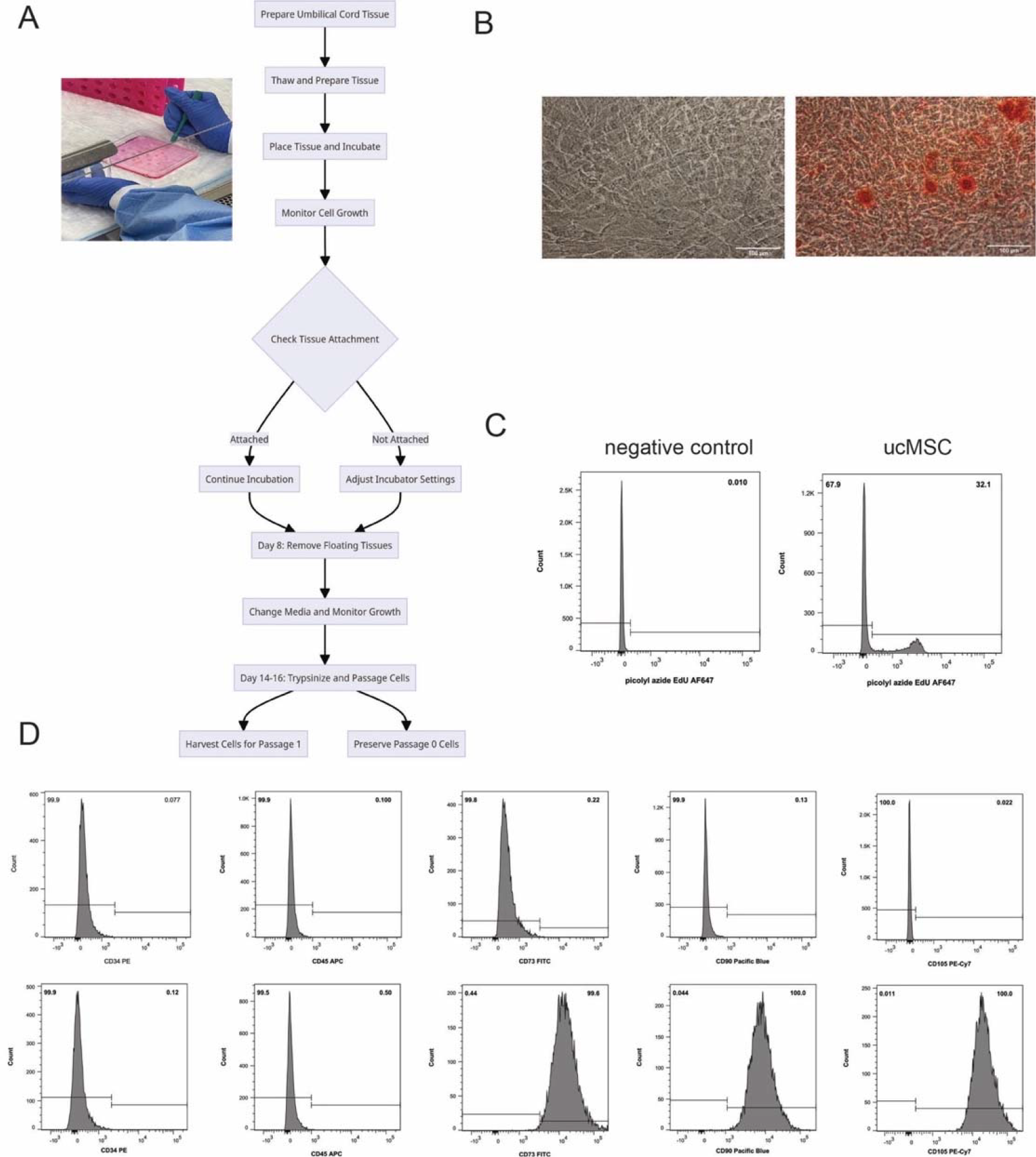
Derivation and Characterization of Mesenchymal Stem Cells from Umbilical Cord Tissue. A. MSC Isolation from Umbilical Cord Tissue (ucMSCs): The methodology employed in extracting MSCs from umbilical cord tissue with detailed steps from tissue processing to the establishment of cell culture conditions and subsequent cell expansion strategies. B. Morphological Assessment and Osteogenic Differentiation: The cellular morphology of ucMSCs derived from a healthy donor alongside their osteogenic differentiation capabilities on day 10, as evidenced by Alizarin-Red staining to identify calcium deposits indicative of osteogenesis. C. Proliferation Capacity of ucMSCs: The 2-hour EdU proliferation assay highlighting the ucMSCs’ significant proliferative ability. D. Surface Marker Characterization of ucMSCs: ucMSCs showing the positive expression of specific MSC surface markers (CD73, CD90, and CD105) and the absence of expression for hematopoietic markers (CD34 and CD45), underscoring their MSC identity.

### Characterization of ucMSC from frozen cord tissues

These MSCs, referred to as ucMSCs, were subjected to an osteogenic differentiation medium, under which they were successfully differentiated into mature osteoblasts. The mineralized extracellular calcium deposits were visualized by Alizarin-Red staining (Figure 1B). The staining results confirmed the presence of calcium deposits, underscoring the differentiation of ucMSCs into osteoblasts. In addition to their differentiation capacity, ucMSCs exhibited a high proliferative capacity, showing 32.1% EdU incorporation (Figure 1C), making them highly suitable for reprogramming where cell proliferation is desirable. To further characterize the isolated ucMSCs, fluorescence-activated cell sorting (FACS) analysis showed that ucMSCs strongly expressed MSC markers, such as CD73, CD90, and CD105, establishing they are MSCs. Conversely, the ucMSCs were negative for the blood cell markers CD34 and CD45 (Figure 1D). This comprehensive characterization confirms the mesenchymal stem cell identity of ucMSCs.

### Chemical reprogramming of ucMSC to human chemical induced iPSC (hCiPSC)

To initiate the chemical reprogramming of ucMSCs, the cells were seeded at a density of 1–1.5 × 10^4 cells per well in a 12-well plate with DMEM medium supplemented with 15% FBS (Figure 2A). Cells were treated with stage I induction medium the next day and immediately maintained in a 5% O_2_ hypoxia incubator. Cells were cultured in stage I induction medium under hypoxia condition for 8 days. By days 4-6 of stage I, epithelial-like cells began to emerge, achieving approximately 80% confluence. On day 8, cells were switched to stage II induction medium in regular O_2_ (21%) incubator for the rest of process. The emergence of multilayered cell colonies was observed within 8-12 days during stage II. The colonies were expanded rapidly and were cultured with stage III medium for another 12 days. VPA was included for the first four days of the induction process in stage IV medium and the appearance of primary human chemical induced iPSC (hCiPSC) colonies were observed after 6-8 days of culture. Primary colonies from ucMSCs were dissociated and replaced for expansion. Compact hCiPSC colonies were manually picked and mechanically dissected into small clumps as typical iPSC culture (Figure 2B).

**Figure 2.**
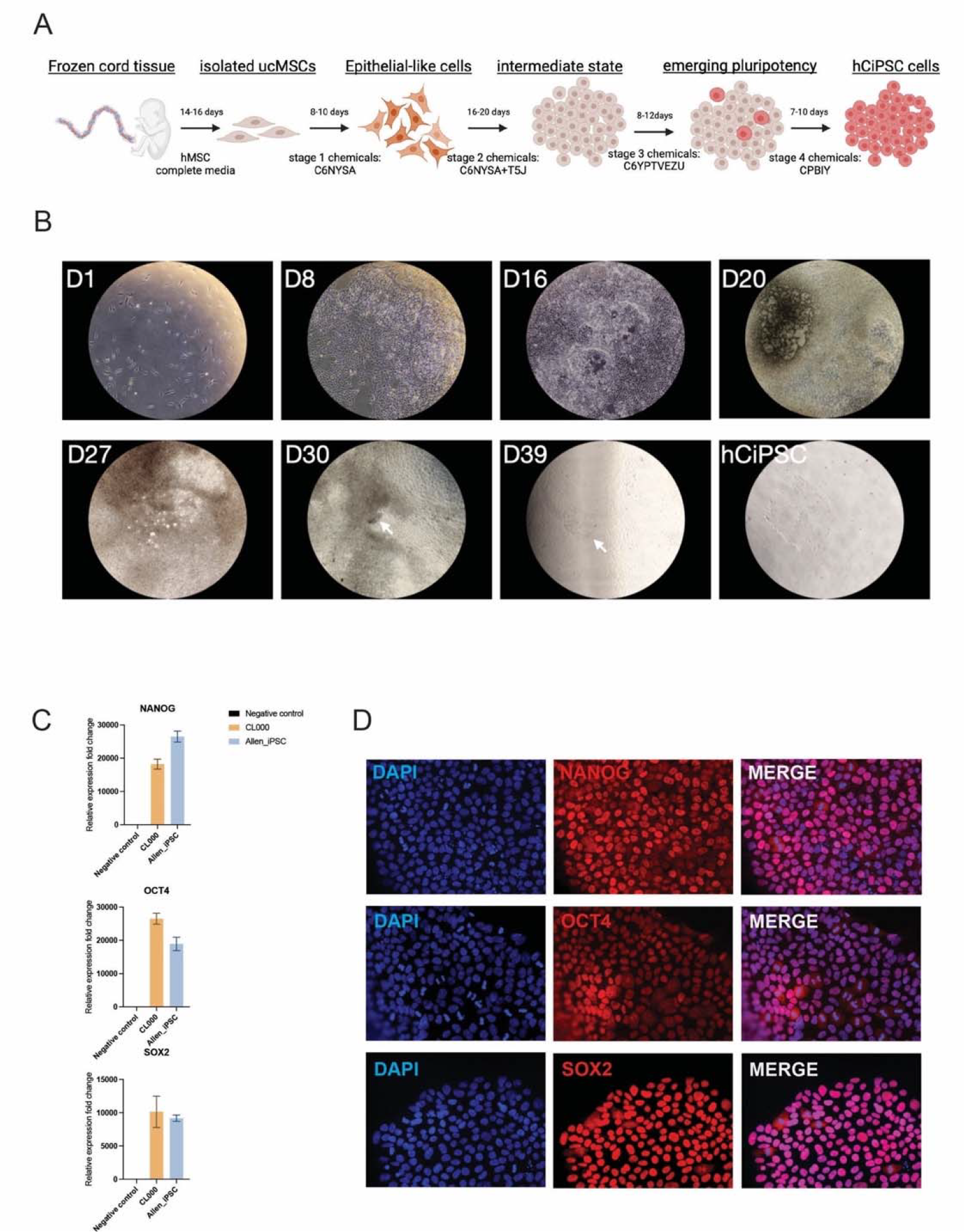
Chemical Reprogramming of ucMSCs to iPSCs. A. Reprogramming Protocol: The specific chemical reprogramming process to transform MSCs from frozen umbilical cord tissue into hCiPSCs, highlighting the different media for each stage and the duration of treatments throughout the process. B. Morphological Transformations: A series of cell images capturing key stages of cellular morphology changes during the chemical reprogramming process. Arrows indicating the emergence of primary hCiPSC colonies. C. Expression level of Pluripotency Genes in hCiPSCs: Comparison of the expression levels of crucial pluripotency genes (Nanog, OCT4, SOX2) in hCiPSCs derived from ucMSCs using qPCR, with human iPSCs from the Allen Institute serving as a positive benchmark and ucMSCs as a negative control. D. Verification of Pluripotency Markers: Immunostaining showing the high-level expression of key pluripotency markers (OCT4, SOX2, NANOG) in hCiPSCs originating from ucMSCs, affirming their pluripotent status.

### Characterization of hCiPSC from ucMSC

To assess the pluripotent characteristics of hCiPSCs generated from ucMSCs, we examined the expression levels of key pluripotency genes, namely NANOG, OCT4, and SOX2. For comparison, we employed a hiPSC line from the Allen Institute as a positive control and ucMSCs as the baseline negative control. Quantitative real-time PCR (qPCR) analysis showed comparable expression level of NANOG, OCT4, and SOX2 between our hCiPSCs and the iPSCs from Allen Institute, which were undetectable in ucMSCs (Figure 2C). Furthermore, Immunofluorescence staining revealed robust protein expression of these pluripotency markers in hCiPSCs, reinforcing their pluripotent status (Figure 2D).

## DISCUSSION

In this study, we used a small molecule chemical reprogramming strategy to generate human pluripotent stem cells from frozen umbilical cord tissues. Human somatic cells are resistant to chemical stimuli by exhibiting a stable epigenome and low plasticity, the restricted human epigenetic landscape can be unlocked into a plastic state small molecules, allowing the release of small molecules into human cells and reprogramming of somatic cells into induced pluripotent stem cells^1^.

Currently the most commonly used method is Sendai virus reprogramming by introducing the reprogramming factors OCT4, SOX2, KLF4, and CMYC (OKSM) ^2,9^. However, the continued expression of exogenous Yamanaka factors via Sendai virus over multiple passages poses significant teratoma risks for clinical use^10^. Notably, c-Myc is one of the most frequently mutated genes in human cancer, and previous studies have shown that chimeric mice generated from iPSC by inducing retrovirus-mediated transfection of four reprogramming factors frequently develop tumors ^11^.

In contrast, chemical reprogramming offers a safer and more controlled approach by eliminating the need for introducing exogenous factors like viral vectors or genetic modifications. Chemical reprogramming involves the use of small molecules to induce cellular dedifferentiation and pluripotency, without permanently altering the cell’s genome or introducing foreign elements. Small molecules can be readily synthesized, optimized, and standardized, allowing for reproducible and cost-effective protocols that can be easily adapted for various cell types and applications. This scalability is crucial for the widespread implementation of cell-based therapies and regenerative medicine approaches. Previous studies have found that mouse somatic cells can be reprogrammed into iPSCs via small molecule compounds^12^. The recent success of chemical reprogramming from human adipose derived mesenchymal stem cells is a paradigm shift in reprogramming technology.

Various types of somatic cells have been shown to be reprogrammed to generate induced pluripotent stem cells. Cell proliferation has been shown to affect reprogramming efficiency. In contrast, reprogramming efficiency is negatively associated with donor age, and aged cells retain their DNA mutations during reprogramming ^8^. In this regard, umbilical cord cells represent the youngest cells that can be used to create iPSCs ^13^. Umbilical cord tissue MSCs can be easily isolated, have the same advantages as umbilical cord blood cells, and are simple to operate ^14^. In this study, we found that MSCs can be efficiently isolated from frozen umbilical cord tissue and expressed all MSC cell surface markers. They are highly proliferative and can be passaged long term to yield millions of cells during the initial plating as reported^15^. Furthermore, because they are young cells from birth, these cells are absent of environmental risk factors or aging caused somatic mutations, serving as a perfect source for studying familial mutation associated disease, such as congenital heart disease.

Leveraging the advantage of umbilical cord MSCs and chemical reprogramming method, we have successfully generated human iPSC from MSCs isolated from frozen umbilical cord tissue using chemical reprogramming. This approach provides a promising avenue for expanding cell sources for iPSC reprogramming and advance regenerative medicine applications, particularly in cardiovascular diseases.

## MATERIALS AND METHODS

### ucMSC isolation and culture

ucMSCs were isolated from frozen umbilical cord tissue as described above following a published protocol^16^. Complete MSC medium were prepared with MEM Alpha (1X) + GlutaMAX™ -1 (GIBCO, 32561037), PLTMax GMP Clinical Grade Supplement (Mill Creek Life Sciences, PLTMax27GMP), and Heparin Solution (0.2%) (STEMCELL Technologies, 07980). CELLstart™ Humanized Substrate for Cell Culture (GIBCO, A10142-01) was used for tissue segment attachment and ucMSC expansion.

### Chemical reprogramming medium Stage I Induction Medium

KnockOut DMEM (Gibco, 10829018) supplemented with:10% KnockOut Serum Replacement (KSR) (Gibco, 10828028), 10% FBS, 1% GlutaMax, 1% NEAA, 0.055 mM 2-mercaptoethanol. Small molecules: 50 μg/ml L-ascorbic acid 2-phosphate (Vc2P) (Sigma-Aldrich, A8960), 5 mM LiCl (Sigma-Aldrich, L4408), 1 mM nicotinamide (NAM) (Sigma-Aldrich, 72340), 2 mg/ml AlbuMax-II (Gibco, 11021045), Small molecules: CHIR999021 (10 μM), 616452 (10 μM), TTNPB (2 μM), SAG (0.5 μM), ABT-869 (1 μM), Rock inhibitor (Y-27632 (2 μM) or Tzv (2 μM)).

### Stage II Induction Medium

KnockOut DMEM supplemented with: 10% KSR, 10% FBS, 1% GlutaMax, 1% NEAA, 0.055 mM 2-mercaptoethanol, 50 μg/ml Vc2p, 5 mM LiCl, 1 mM NAM, 40 ng/ml bFGF (Origene, TP750002).

Small molecules: CHIR99021 (10-12 μM), 616452 (10 μM), TTNPB (2 μM), SAG (0.5 μM), ABT-869 (1 μM), Y27632 (10 μM), JNKIN8 (1 μM), tranylcypromine (10 μM), 5-azacytidine (10 μM), UNC0224 (1 μM), ruxolitinib (1 μM), SGC-CBP30 (2 μM).

### Stage III Induction Medium

KnockOut DMEM supplemented with: 1% N2 supplement (Gibco, 17502-048), 2% B27 supplement (Gibco, 17504-044), 1% GlutaMax, 1% NEAA, 0.055 mM 2-mercaptoethanol, 50 μg/ml Vc2p, 5 mg/ml AlbuMax-II, 20 ng/ml recombinant human heregulin β-1 (HRG) (PeproTech, 100-03). Small molecules: CHIR99021 (1 μM), 616452 (10 μM), Y-27632 (10 μM), PD0325901 (1 μM), tranylcypromine (10 μM), VPA (500 μM), DZNep (0.2 μM), EPZ004777 (5 μM), UNC0379 (1 μM).

### Stage IV Induction Medium

KnockOut DMEM supplemented with: 1% N2 supplement, 2% B27 supplement, 1% GlutaMax, 1% NEAA, 0.055 mM 2-mercaptoethanol, 50 μg/ml Vc2p, 20 ng/ml HRG. Small molecules: CHIR99021 (1 μM), Y-27632 (10 μM), PD0325901 (1 μM), IWP-2 (2 μM), SB590885 (0.5 μM), with VPA (500 μM) included for the first 4 days.

### Derivation and culture of hCiPSCs

Following an 8-12 day treatment in Stage IV conditions, cells were dissociated using Accutase (Millipore) and replated onto Laminin coated plate at 1:12 ratio, with a modified Stage IV medium. This medium consisted of KnockOut DMEM enhanced with 1% N2 and 2% B27 supplements, 1% GlutaMax, 1% NEAA, 1% penicillin–streptomycin, 0.055 mM 2-mercaptoethanol, 50 μg/ml Vc2p, 2 mg/ml AlbuMax-II, along with small molecules including CHIR99021 (1 μM), PD0325901 (0.5 μM), IWP-2 (2 μM), Y-27632 (10 μM), HRG (20 ng/ml), and bFGF (100 ng/ml from Peprotech). Culture medium was changed daily. Within 7 days, hCiPS cell colonies started to form. After 4-6 days growing, these colonies were mechanically cut into smaller clusters, and transferred to plates coated with Matrigel (Corning, 354248) in StemMACS™ iPS-Brew XF medium (Miltenyi Biotec, 130-104-368) supplemented with Y-27632 (10 μM). After 24 hours cells were switch into iPS-Brew XF medium without Y-27632.

### MSC Osteogenic Differentiation

ucMSCs were cultured in Mesenchymal Stem Cell Osteogenic Differentiation Medium (PromoCell, C-28013) for osteogenic differentiation. After 10 days of differentiation, cells were washed twice with PBS and fixed in 4% paraformaldehyde for 30 minutes. Cells were then stained by adding sufficient Alizarin-Red Staining Solution (Sigma-Aldrich, TMS-008-C) following manufacturer’s instruction.

### Fluorescence-activated Cell Sorting (FACS) analysis for Cell Surface Staining and MSC markers

Proliferation assay: EdU (10 μM) was added to the ucMSC culture medium 2hr before harvest. The cells were then trypsinized and fixed in 4% formaldehyde. EdU incorporation was determined with the Click-iT™ EdU Alexa Fluor™ 647 Flow Cytometry Assay Kit (Invitrogen, C10419) according to the manufacturer’s instructions.

MSC markers: A total of 5×10^5^ cells were resuspended and incubated with fluorescence-conjugated antibody mixture (CD34-PE; CD45-APC; CD73-FITC; CD90-Pacific Blue; CD105-PE-Cy7) for 20 min at room temperature. Next, cells were incubated with biotinylated secondary antibodies for 30 min on ice followed FACS buffer wash for 3 times. The fluorescence intensity of the cells was measured using a flow cytometer.

### Quantitative RT-PCR analysis

Total RNA was prepared from the hCiPSC using the RNeasy Plus Mini Kit (Qiagen). Reverse transcription was performed using the High-Capacity cDNA Reverse Transcription Kit (Life Technologies). Quantitative RT-PCR was carried out using CFX96 real-time PCR detection system (Bio-Rad) with TaqMan Universal PCR Master Mix (Applied Biosystems). The relative mRNA expression of the targeted genes was normalized to the GADPH as endogenous control. Gene expression was calculated using 2–ΔΔCt method. Human iPSC line (AICS-0060-027iPSC from Stem cell) purchased from Allen Institute was used as positive control.

### Immunofluorescence staining

Immunofluorescence staining was performed to analyze OCT4, SOX2, NANOG protein expression. Cells fixed using 4% paraformaldehyde for 2d and washed with 0.1% Triton X-100 (Sigma-Aldrich). Blocking was done using 2.5% donkey serum (Sigma-Aldrich) and then incubated with the primary antibodies (Cell signaling Technology, StemLight™ Pluripotency Antibody Kit #9656) in 2.5% donkey serum–PBS for overnight at 4°C. On the next day, after three washes with PBS, the cells were further incubated with secondary antibodies (Thermo Fisher, Alexa Fluro 488 and 594) for 1 hour at room temperature. Next, cells were washed with PBS and mounted with Slow Fade Gold Antifade Reagent with 4,6-diamidino-2-phenylindole (DAPI; Life Technologies) before imaging. All immunofluorescent images were captured using a Nikon Eclipse 80i fluorescence microscope.

## REFERENCES

1. Guan J, Wang G, Wang J, Zhang Z, Fu Y, Cheng L, Meng G, Lyu Y, Zhu J, Li Y, et al. Chemical reprogramming of human somatic cells to pluripotent stem cells. Nature. 2022;605:325–331. doi: 10.1038/s41586-022-04593-5

2. Kunitomi A, Hirohata R, Arreola V, Osawa M, Kato TM, Nomura M, Kawaguchi J, Hara H, Kusano K, Takashima Y, et al. Improved Sendai viral system for reprogramming to naive pluripotency. Cell Rep Methods. 2022;2:100317. doi: 10.1016/j.crmeth.2022.100317

3. Liuyang S, Wang G, Wang Y, He H, Lyu Y, Cheng L, Yang Z, Guan J, Fu Y, Zhu J, et al. Highly efficient and rapid generation of human pluripotent stem cells by chemical reprogramming. Cell Stem Cell. 2023;30:450–459 e459. doi: 10.1016/j.stem.2023.02.008

4. Lange L, Esteban MA, Schambach A. Back to pluripotency: fully chemically induced reboot of human somatic cells. Signal Transduct Target Ther. 2022;7:244. doi: 10.1038/s41392-022-01109-5

5. Nagamura-Inoue T, Mukai T. Umbilical Cord is a Rich Source of Mesenchymal Stromal Cells for Cell Therapy. Curr Stem Cell Res Ther. 2016;11:634–642. doi: 10.2174/1574888x10666151026115017

6. Beeravolu N, McKee C, Alamri A, Mikhael S, Brown C, Perez-Cruet M, Chaudhry GR. Isolation and Characterization of Mesenchymal Stromal Cells from Human Umbilical Cord and Fetal Placenta. J Vis Exp. 2017. doi: 10.3791/55224

7. Wang Q, Wang Y, Chang C, Ma F, Peng D, Yang S, An Y, Deng Q, Wang Q, Gao F, et al. Comparative analysis of mesenchymal stem/stromal cells derived from human induced pluripotent stem cells and the cognate umbilical cord mesenchymal stem/stromal cells. Heliyon. 2023;9:e12683. doi: 10.1016/j.heliyon.2022.e12683

8. Trokovic R, Weltner J, Noisa P, Raivio T, Otonkoski T. Combined negative effect of donor age and time in culture on the reprogramming efficiency into induced pluripotent stem cells. Stem Cell Res. 2015;15:254–262. doi: 10.1016/j.scr.2015.06.001

9. Takahashi K, Tanabe K, Ohnuki M, Narita M, Ichisaka T, Tomoda K, Yamanaka S. Induction of pluripotent stem cells from adult human fibroblasts by defined factors. Cell. 2007;131:861–872. doi: 10.1016/j.cell.2007.11.019

10. Angeles ADL, Hug CB, Gladyshev VN, Church GM, Velychko S. Sendai virus persistence questions the transient naive reprogramming method for iPSC generation. bioRxiv. 2024:2024.2003.2007.583804. doi: 10.1101/2024.03.07.583804

11. Okita K, Ichisaka T, Yamanaka S. Generation of germline-competent induced pluripotent stem cells. Nature. 2007;448:313–317. doi: 10.1038/nature05934

12. Hou P, Li Y, Zhang X, Liu C, Guan J, Li H, Zhao T, Ye J, Yang W, Liu K, et al. Pluripotent stem cells induced from mouse somatic cells by small-molecule compounds. Science. 2013;341:651–654. doi: 10.1126/science.1239278

13. Zhou H, Rao MS. Can cord blood banks transform into induced pluripotent stem cell banks? Cytotherapy. 2015;17:756–764. doi: 10.1016/j.jcyt.2015.02.008

14. Mohamed A, Chow T, Whiteley J, Fantin A, Sorra K, Hicks R, Rogers IM. Umbilical Cord Tissue as a Source of Young Cells for the Derivation of Induced Pluripotent Stem Cells Using Non-Integrating Episomal Vectors and Feeder-Free Conditions. Cells. 2020;10. doi: 10.3390/cells10010049

15. Raileanu VN, Whiteley J, Chow T, Kollara A, Mohamed A, Keating A, Rogers IM. Banking Mesenchymal Stromal Cells from Umbilical Cord Tissue: Large Sample Size Analysis Reveals Consistency Between Donors. Stem Cells Transl Med. 2019;8:1041–1054. doi: 10.1002/sctm.19-0022

16. Skiles ML, Marzan AJ, Brown KS, Shamonki JM. Comparison of umbilical cord tissue-derived mesenchymal stromal cells isolated from cryopreserved material and extracted by explantation and digestion methods utilizing a split manufacturing model. Cytotherapy. 2020;22:581–591. doi: 10.1016/j.jcyt.2020.06.002

